# Distinct visual pathways of threat retrieval in fear-conditioned faces

**DOI:** 10.64898/2026.03.13.711521

**Authors:** Enya M. Weidner, Marlene Götze, Adrian Taday, Johanna Kissler

**Author notes:** **Contact information:** Enya Weidner, Marlene Götze, Adrian Taday, Johanna Kissler.

## Abstract

Numerous studies have demonstrated rapid (< 100 ms) visuo-cortical differentiation of threat-associated faces. This may be due to low-spatial frequency (LSF) visual information originating from magnocellular pathways. Yet it remains unclear whether potentially magnocellular fear signals extend beyond evolutionarily prepared emotional faces and whether they are subject to short-term neuroplasticity. If so, spatial frequency characteristics should modulate processing of faces with newly acquired threat-relevance. Furthermore, it is unknown whether sub-bands of the visual spectrum are associated with autonomic arousal. Using a differential fear-conditioning paradigm, this study tested whether early visual attentional capture, indicated by the P1 event-related potential component, prioritizes LSF information of threat-associated faces with neutral expressions. Additionally, it was tested whether such effects would be paralleled by threat differentiation in the skin conductance response (SCR). For contingency aware participants, stimulus ratings confirmed successful fear conditioning and participants showed a selective left-hemispheric enhancement of the P1 in response to LSF threat-faces. By contrast, CS differentiation in the SCR was not modulated by spatial frequencies but by stimulus duration, with longer CS presentations resulting in larger SCR to threat compared to neutral faces. For contingency unaware participants, trial-by-trial amplitudes of P1 and SCR were positively correlated. Data support the notion that magnocellular-cortical pathways adapt quickly to novel threat-associations and facilitate rapid threat retrieval even for perceptually neutral faces. However, at least in the short term, these signals do not necessarily associate with anticipatory arousal in SCR.

**Impact statement:** Our electroencephalography (EEG) study provides evidence for distinct contributions of subcortical signals during early visual perception of fear-conditioned faces (P1 event-related potential) but not autonomic arousal (skin conductance response). Instead, skin conductance responses reflected conscious anticipatory arousal irrespective of the visual pathway. Together, these results reveal parallel but dissociable mechanisms of fear perception that are differentially sensitive to visual properties of threat-associated faces.

## Introduction

Rapid (< 100 ms) visuocortical processing advantages of threat-relevant faces have often been interpreted as evidence for a neural shortcut of fear (Carretié et al., 2017; Garvert et al., 2014; LeDoux, 2007; Morris et al., 1999). Presumably, coarse low spatial frequency (LSF) information transmitted by magnocellular subdivisions of the lateral geniculate nucleus suffices to generate amplified electrophysiological responses to threat-related faces (Méndez-Bértolo et al., 2016; Vuilleumier et al., 2003; Winston et al., 2003). This would bypass slower visuocortical, parvocellular processing of high spatial frequencies (HSF) (Crook et al., 1988; Maunsell & Gibson, 1992; McFadyen et al., 2019; Purpura et al., 1988; Schiller & Malpeli, 1978). Still, the relative contribution of these pathways to threat differentiation in early visual attention remains debated (Boatman & Kim, 2006; Li & Keil, 2023; McFadyen et al., 2017). In fact, a common marker for early (∼90 – 130 ms) visual attention, the P1 event-related potential (ERP) component in surface electroencephalogram (EEG) studies, has shown unstable threat-sensitivity across studies (Schindler & Bublatzky, 2020) and low-level visual properties have been suggested as confounds (Schindler, Bruchmann, et al., 2021). Understanding the neural pathways that underlie threat-detection in the P1 might help to align these inconsistencies.

Favoring the idea of magnocellular contributions, the P1 has been shown amplified in response to LSF relative to HSF images (Cesarei et al., 2013) and faces (Nakashima et al., 2008). Crucially, Bruchmann and colleagues (2020) found P1 amplitudes to differentiate fearful from neutral faces only when LSF spectra of fearful faces were preserved. What remains uncertain is whether LSF fearful faces are so readily attended to in early visual processing because their stereotypical perceptual properties and configuration in LSF (Barrett, 2018; Bruchmann et al., 2020; Goffaux et al., 2005; Tipples, 2007) enhances salience more strongly than the actual concept of threat. Correspondingly, fear-neutral differentiation in the P1 was found preserved even for scrambled faces (Schindler, Wolf, et al., 2021). By contrast, Yu et al. (2023) argue that conceptual threat relevance but not perceptual information modulates the P1: Although observed for upright faces, they found no P1 enhancement for inverted angry versus neutral faces containing the same physical information. However, the different sets of facial expressions used across studies might limit their comparability (N’Diaye et al., 2009; Springer et al., 2007). Fear conditioning can be used to robustly manipulate conceptual threat-relevance without confounding perceptual differences (Lonsdorf & Kalisch, 2011; Stolarova et al., 2006). Utilizing this paradigm, many studies have already reported early-onset (∼ 100 ms) increases of occipito-temporal electrophysiological responses, including the P1, to neutral faces associated with an aversive outcome (Bruchmann et al., 2023; Junghöfer et al., 2017; Linton & Levita, 2021; Moratti et al., 2006; Roesmann et al., 2020; Sperl et al., 2021; Steinberg et al., 2012). Early effects of fear conditioning have also been found to spatially extend to frontal and parietal regions (Hintze et al., 2014; Petro et al., 2017; Rehbein et al., 2014; Rehbein et al., 2015; Roesmann et al., 2022; Steinberg et al., 2012), suggesting a parallel engagement of multiple neural pathways during early threat perception. In fact, studies have proposed hemispheric differences in processing of spatial frequencies (Kitterle et al., 1990; Peyrin et al., 2003; Rebaï et al., 1998), with particularly magnocellular pathways being left-dominant, while face processing commonly associated with right-hemispheric systems (Fox et al., 2009; Kanwisher et al., 1997; Kanwisher & Yovel, 2006). To date, the lateralization of spatial frequency (SF) specific pathways during facial threat perception remains unclear.

Despite this body of research, to the best of our knowledge, fear conditioning has not been utilized to compare magnocellular and parvocellular contributions to the processing of newly acquired facial threat while controlling perceptual confounds. In addition, it remains largely unexplored whether distinct visual pathways of threat processing are reflected in autonomic arousal. Prior functional magnetic resonance imaging (fMRI) studies have repeatedly shown that the skin conductance response (SCR) tracks fear learning in parallel to sensory, frontal, and medial temporal areas (Büchel & Dolan, 2000; Büchel et al., 1999; Büchel et al., 1998). Similarly, many EEG studies demonstrated that enhanced visuocortical responses to fear-conditioned stimuli co-occur with increasing SCR (Antov et al., 2020; Gruss & Keil, 2019; Pastor et al., 2015; Stegmann et al., 2024). Highlighting the role of SF information for fear retrieval in the SCR, Chen et al. (2023) show that fear-conditioned SCR to perceptually neutral, abstract shapes was partly mediated by LSF for unconsciously evaluated fear-conditioned stimuli. Hence, autonomic arousal might be sensitive to specific visual features. But given that the SCR is a slow, long-latency signal that might well be modulated by factors beyond early visual attention (Sevenster et al., 2014), its relationship to SF-specific effects in the P1 during threat processing remains unknown.

Against this background, we investigated whether processing of fear-conditioned faces in the P1 and SCR is facilitated by specific SF bands of the visual spectrum. In our paradigm, greyscale photographs of neutral facial expressions containing a broad-band frequency spectrum were either fear- or safety-conditioned, mimicking typical fear acquisition settings (Junghöfer et al., 2017; Moratti et al., 2006; Roesmann et al., 2020; Sperl et al., 2021; Steinberg et al., 2012). SF-specific threat retrieval was subsequently probed by presenting LSF- and HSF-filtered versions of the conditioned faces while EEG was recorded. Our preregistered hypotheses (https://osf.io/5kbmn/) state that the P1 (Bruchmann et al., 2020) and SCR (Chen et al., 2023) should show highest amplitudes for threat-associated faces that are shown in LSF. Furthermore, stimulus presentation time was varied randomly (100/1000 ms). Shorter presentation times could facilitate magnocellular signaling by suppressing saccadic activity (Ross et al., 1996) but also limit sustained, conscious differentiation of facial identities (Tanskanen et al., 2007) that might specifically affect stimulus discrimination in the SCR (Baeuchl et al., 2019; Sevenster et al., 2014). Therefore, we expected stronger SCR threat differentiation for longer stimulus durations (Sevenster et al., 2014; Tanskanen et al., 2007). We also explored contingency awareness as a potential mediator of CS-differentiation, as, at least for the SCR, contingency awareness is assumed to be a prerequisite for threat-specific responses (Baeuchl et al., 2019).

## Method

### Sample

The initial sample consisted of 77 participants. No participant reported any psychiatric or neurological disorders, nor the use of drugs in the two weeks prior to testing. Data from four participants were excluded due to poor recording quality and five subjects had excessive artefactual EEG signals. The final sample for the EEG analysis comprised 68 participants (age: M = 25.56 years, SD = 7.27 years; 71.01 % female), of whom 58 reported to be right-handed. Due to technical difficulties with the SCR recordings, valid SCR data were available for only 40 participants. Consequently, we conducted additional EEG analyses including only participants for whom both measures were available, which did not alter the pattern of results (see Tables S6 to S8). A priori power analysis was not conducted because the study employed nonparametric permutation-based inference of multilevel data (Manley, 2020; Noguchi et al., 2021; Raychaudhuri, 2008). Therefore, data distribution was not estimated beforehand. Post-hoc power analyses of the decisive effects are reported in the results section. Participants received either €10 or one study credit per hour. The study was conducted in accordance with the Declaration of Helsinki and was approved by the Ethics committee of Bielefeld University (EUB-2024_245), following the ethics guidelines of the German Psychological Association (DGPs).

### Stimuli

We presented neutral facial expressions of four Caucasian male identities (conditioned stimulus, [CS]) from the FACES database (Ebner et al., 2010) with previous permission. Faces were rendered equiluminescent to each other. The isolated, elliptically cropped faces without visible hair, shown in greyscale, were centrally presented on a black screen, measuring 8 cm in width and 12 cm in height (9° 47’ 0.89’’ degree visual angle [centered], Fig. 1a). Stimuli were presented on a DELL Precision 3650 computer with an ASUS VG24BQE Full HD screen using PsychoPy (version 2024.1.5). The aversive unconditioned stimulus (US) was operationalized as a 1 second white noise burst (90 – 95 dB) that was presented binaurally via stereo Creative 265 loudspeakers. The volume was configured with an audiometer. During the acquisition phase, original broadband versions of the faces were presented. For the retrieval phase, CS were spatially filtered and presented either as LSF (≤ 10 cycles/image; ∼0.034 cycles/pixel horizontally, ∼0.023 cycles/pixel vertically) or HSF images (≥ 35 cycles/image; ∼0.120 cycles/pixel horizontally, ∼0.080 cycles/pixel vertically). SF characteristics were chosen based on reported SF sensitivities of magno- and parvocellular pathways (Lynch et al., 1992; Merigan & Katz, 1990) as well as previous studies on SF-specific face perception (Bruchmann et al., 2020; Méndez-Bértolo et al., 2016) and extracted using custom MATLAB scripts (version 2019b, The MathWorks Inc.).

**Figure 1.**
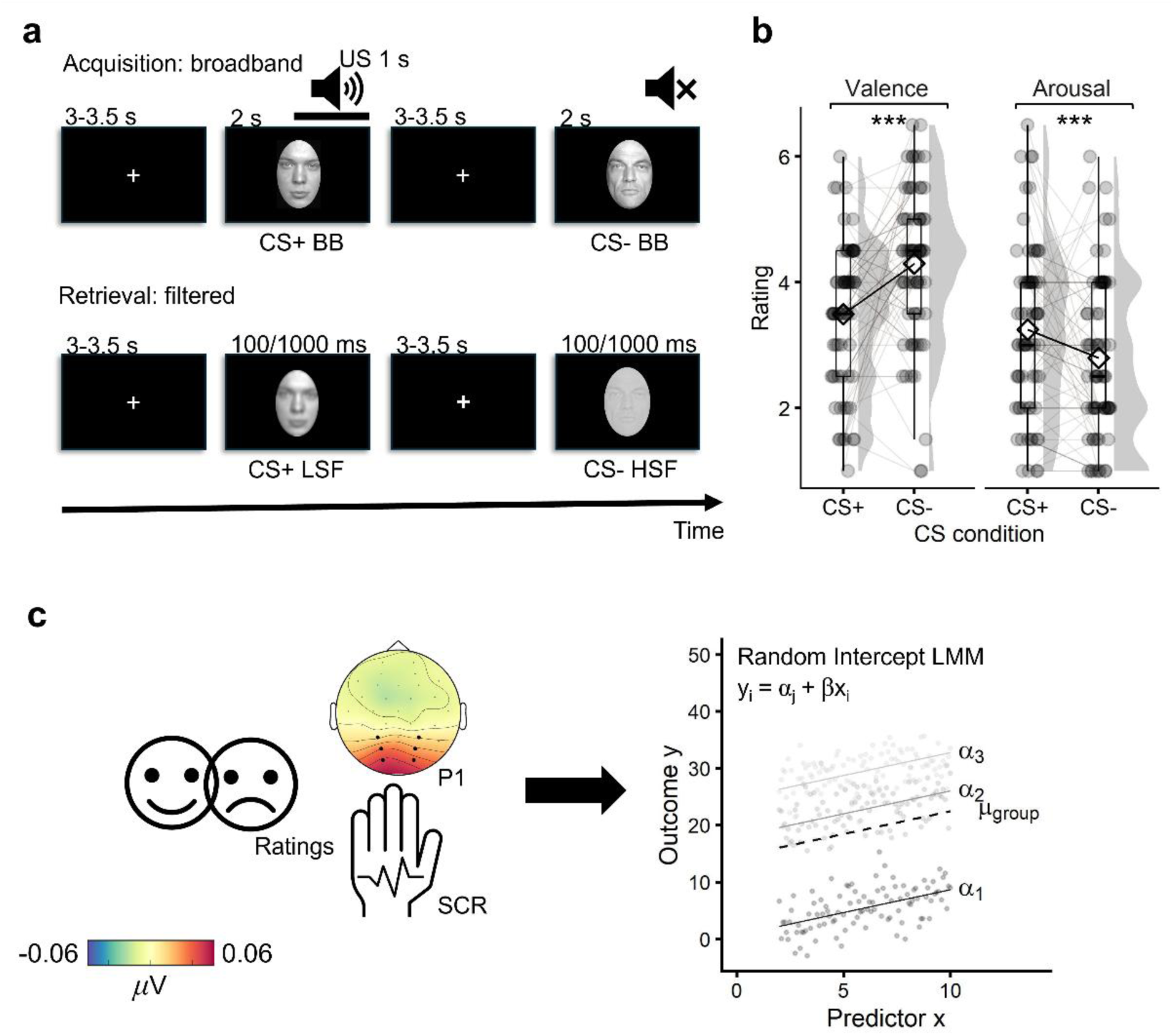
Experimental paradigm and validation of threat-induction. **a** Exemplary trial and dependent variables in the acquisition and the retrieval phase. Faces are derived from the FACES database (Ebner et al., 2010) with previous permission. Please note that the faces shown were not used in the experiment but selected to comply with publishing guidelines. **b** Stimulus ratings of valence and arousal. Boxplots show median and interquartile range; raincloud plots display single-subject means per *CS* condition along with the smoothed distribution density; diamonds indicate the overall condition mean. Line plots depict data trends. The black line shows the mean across all subjects; grey lines show single-subject trends. Brackets mark significant effects of *CS* on ratings (\*\*\**p* ≤ .001). **c** Dependent variables and illustration of the analysis approach. The topography shows mean ERP amplitudes in the P1 time-window. The P1 was extracted from 90 – 130 ms after stimulus onset from the channels marked in black. SCR was measured as the first-interval response in a 900 – 4500 ms trough-to-peak window after stimulus onset. A linear mixed model (hierarchical regression) was implemented for all measures of interest. *Abbreviations*: BΒ = Broadband, CS = conditioned stimulus, LMM = linear mixed model, LSF = low spatial frequency, HSF = high spatial frequency, SCR = skin conductance response, US = unconditioned stimulus.

### Procedure

Before the experiment, participants were asked to give their informed consent. Demographic as well as psychiatric variables were documented. Psychiatric assessments included the Becks-Depression-Inventory (BDI; *M* = 6.71, *SD* = 6.70; Beck et al., 1996), the State-Trait-Anxiety Inventory (STAI; *M*_State_ = 32.89, *SD*_State_ = 6.15, *M*_Trait_ = 37.55, *SD*_Trait_ = 8.25; Spielberger, 1970), and the Harvard Trauma Questionnaire (HTQ; *M*_self-experienced_ = 0.81 event, *SD*_self-experienced_ = 1.72 events; Kleijn et al., 2001). Then, the EEG headcap, electrodes, and electrolyte gel were applied. Before the experiment started, participants were able to individually adjust US volume between 90 and 95 dB (Bierwirth et al., 2021; Sperl et al., 2016) so that the noise was highly uncomfortable but not painful (Mueller et al., 2014). Afterwards, the experiment started with a habituation phase, consisting of 5 presentations of the US and subsequently 4 presentations of each CS. Then, the acquisition phase started with the pseudo-randomized presentation of 20 unfiltered presentations of each CS. Here, two out of four identities were intermittently paired with the US in 50 % of occurrences (CS+) while the remaining two were not paired with any sound (CS-). Partial reinforcement was chosen to avoid habituation to the US and to delay extinction in the retrieval phase described below (Lonsdorf et al., 2017). The assignment of face identities to the CS conditions was randomized across participants. All CS were presented for 2000 ms, the US was presented 1000 ms after CS+ onset. The interstimulus interval consisted of a white fixation cross with a variable presentation time of 3000 – 3500 ms to reduce stimulus onset predictability. Overall, the first acquisition phase consisted of 80 trials. In the ensuing retrieval phase, CS were presented in either LSF or HSF. The presentation duration was randomly varied so that approximately 50 % of stimuli in each *CS* × *SF* condition were presented for 100 ms, and 50 % for 1000 ms. Each *CS* × *SF* × *Presentation time* cell consisted of 20 trials, leading to a total of 160 trials per retrieval block. To avoid early extinction in the retrieval phase (Bierwirth et al., 2023; Lissek et al., 2008; Sperl et al., 2019), CS+ were still randomly paired with an US after 200 ms with a 25 % chance. An exemplary trial run is illustrated in Fig. 1a.

Acquisition and retrieval blocks were repeated two more times. The acquisition blocks were reduced to 40 trials per block (10 trials/identity). Trial count in the retrieval block was equal across blocks. Blocks were separated by self-paced breaks. Across all phases, consecutive presentations of the same identity were constrained to a maximum of four trials. After the last retrieval phase, we included a brief extinction run, presenting all CS identities and no US for four times each (16 trials in total). Afterwards, participants were asked to rate both arousal and valence of all four CS identities on a 7-point Likert scale (Bradley & Lang, 1994). Furthermore, we assessed contingency awareness and US expectancy during the retrieval phase with questionnaires. Contingency awareness was considered given if participants correctly indicated the regularity in the occurrence of the US or described the CS+ faces as aversive and CS- faces as neutral/non-aversive. 49.28 % of participants correctly identified the CS-US contingencies. For US expectancy, 44.93 % of participants reported that they expected the US for up to 10 trials in the retrieval phase, 40.58 % expected it for up to 20 trials and 11.59 % expected it for more than 20 trials. In total, the experiment consisted of 672 CS presentations and took about 60 minutes.

### Dependent measures

#### Ratings

Ratings of arousal and valence were given for all four CS identities on a 7-point Likert scale (Bradley & Lang, 1994). For valence, higher scores (maximum = 7) indicated a more positive valence. For arousal, higher scores (maximum = 7) indicated higher physiological arousal. No information regarding the US contingency was given when ratings were applied.

#### EEG

The EEG was continuously recorded from 32 active BioSemi Ag-AgCl electrodes using the ActiveTwo system and ActiView software (http://www.biosemi.com/). The recording was referenced to the Cz electrode, with the sampling rate set to 1024 Hz. Impedances were kept below 25 kΩ, in line with system recommendations. NaCl contact electrolyte gel was used to improve surface conductivity. Pre-processing of offline data was performed using the FieldTrip toolbox (Version 20240414, Oostenveld et al., 2011) in MATLAB (Version 2019b, The MathWorks Inc.). Continuous offline data were re-referenced to the common average of all EEG channels. A high-pass Butterworth first-order zero-phase filter of 0.1 Hz and a low-pass first-order zero-phase Butterworth filter of 40 Hz were applied to ensure accurate detection of sharp peak components like the P1 (Widmann et al., 2015). Filtered data were cut into segments of -2000 ms to 2000 ms relative to stimulus onset and subsequently down-sampled to 250 Hz. Baseline-correction used 200 ms before stimulus onset. Bad channels were identified using a *z*-value based threshold and temporarily removed before independent component analysis (ICA). Datasets with > 10 bad channels were excluded from the analysis. ICA was performed to identify and remove components associated with ocular, muscle, and technical artifacts. On average, 11.23 % of channels were interpolated per particpant. After the ICA, previously excluded channels were interpolated using a weighted neighboring approach. On average, 4.55 % of channels were interpolated. Finally, if single trials exceeded a maximum amplitude range of 250 μV, they were excluded. For statistical analysis, data were averaged across time-points and channel clusters of the P1. In line with previous studies on facial threat differentiation in the P1 (Müller-Bardorff et al., 2018; Schindler, Bruchmann, et al., 2021; Weidner et al., 2025), we scored the component at two symmetrical parieto-occipital electrode clusters (P3, PO3, O1 and P4, PO4, O2, Fig. 1c) from 90-130 ms after stimulus onset.

#### SCR

For the analysis of the SCR, recommendations by Boucsein (2012) were followed. Electrodermal activity was recorded with two Ag-AgCl electrodes placed on the thenar and hypothenar of the non-dominant hand. Skin was pretreated with 70 % ethanol. NaCl contact electrolytes were used in gel form to improve surface conductivity. Data were recorded with the BioSemi EEG amplifier. The sampling rate was 1024 Hz. Offline data were processed with custom scripts in MATLAB and the Fieldtrip toolbox. Continuous offline data were resampled to 10 Hz. A high-pass Butterworth first-order zero-phase filter of 0.1 Hz and a low-pass first-order zero-phase Butterworth filter of 10 Hz were applied. Phasic skin conductivity was extracted in response to each stimulus -500 – 6000 ms after stimulus onset. SCR were manually scored as the trough-to-peak difference by using a custom-made program according to published guidelines (Boucsein et al., 2012) and blind to stimulus type. SCR were required to reach a minimum amplitude of 0.02 mS (Boucsein, 2012; Kuhn et al., 2022). The trough was identified in an onset latency window of 900 – 4500 ms post-stimulus onset and the first peak was identified in a peak detection window (PDW) of maximally 6000 ms post-SCR onset. This corresponds to the so-called “first-interval response” (Kuhn et al., 2022).

CS+ trials with subsequent US presentations were excluded to avoid overlapping responses in the SCR. To ensure comparability between EEG and SCR data, we used the same approach for the EEG analysis. To even out trial counts between CS+ and CS-, we randomly excluded CS+ trials before statistical analysis. No additional trial rejection was done for the SCR, but participants classified as hyper-responders (≥95% of US response amplitudes, *n* = 3) or non-responders (no US responses, *n* = 2) were excluded from the SCR analysis. The number of remaining trials in the SCR analysis was *M* = 39.685 (*SD* = 0.627) trials per condition in the acquisition phase and *M* = 44.728 (*SD* = 4.964) trials per condition in the retrieval phase. Trial counts did not differ between *SF* or *duration* conditions (SF: *F*(1,33) = 1.845, *p* = 0.184; duration: *F*(1,33) = 0.463, *p* = 0.501).

Because of the additional artifact correction, the number of remaining trials in the EEG analysis differed slightly from the SCR analysis. The number of remaining trials was *M* = 39.236 (*SD* = 2.765) trials per condition in the acquisition phase and *M* = 43.272 (*SD* = 5.601) trials per condition in the retrieval phase. Additionally, trial counts did not differ between *SF* or *duration* conditions (SF: *F*(1,67) = 0.467, *p* = 0.497; duration: *F*(1,67) = 0.476, *p* = 0.493).

### Statistical analysis

#### General approach

All statistical analyses were conducted in R (version 2023.03.1, R Core Team, 2021) using dummy-coded hierarchical linear mixed models (LMM, *lme4* v.1.1-34; Fig. 1c). Unstandardized regression coefficients are reported as effect sizes. This method allows direct inferences about the direction of a given effect and interaction due to a priori defined contrasts (Frömer et al., 2018). Significance of fixed-effects Satterthwaite *t*-values was evaluated with non-parametric Monte Carlo permutation tests (Maris & Oostenveld, 2007; for a similar approach, see Weidner et al., 2025). Specifically, within-subject labels were permuted 1000 times, and for each permutation step, a new LMM was fitted. The distribution of permuted *t*-values was then used to compute empirical *p*-values, defined as the proportion of permuted *t*-values greater than or equal to the observed statistic. All tests were two-tailed, and randomization was constrained within participants to preserve subject-level structure. Post-hoc comparisons were performed with the predicted marginal means using *emmeans* (v.1.8.8, Lenth, 2018). With Holm corrections of post-hoc tests, we accounted for the inflation of false positives in multiple testing (Holm, 1979). For the post-hoc tests, Cohen’s *d* is reported as the effect size. It is interpreted as small when *d* = .20, medium when *d* = .50, and large when *d* = .80 (Cohen, 1988, 1992). All data were analyzed based on single-trial data, with *subject* included as random intercepts to account for repeated measures. Post-hoc power estimations of LMM coefficients were conducted with the *simr* package running with 500 simulations (Green & MacLeod, 2016). Post-hoc power was not estimated for null-effects and exploratory analyses. If not stated otherwise, only significant effects are reported in the text. For readability, *t*-values and degrees of freedom derived from LMMs are not reported in the main text but in the corresponding supplementary tables.

#### Analysis models

For the subjective ratings of valence and arousal, ratings per identity were assigned to the CS conditions. Separate models were fit for the two rating measures. Rating analyses were carried out based on the EEG sample (*n* = 68). For electrophysiological responses, we tested whether responses differed between broadband filtered CS+ and CS- during the acquisition phase. These comparisons were based on a one-factorial design, with P1 amplitudes and phasic SCR peaks analyzed as a function of *CS* (CS+ versus CS-, reference level: CS-). For the P1, the second factor *hemisphere* was introduced to test potential lateralization of effects (left versus right, reference level: right).

For the decisive analyses of the retrieval phase, we employed a 2 × 2 repeated-measures design with the fixed-effects *CS* (CS+ versus CS-, reference level: CS-) and *SF* (LSF versus HSF, reference level: HSF). Again, the third factor *hemisphere* (left versus right, reference level: right) was additionally introduced for the P1 analyses. Furthermore, because the long-latency SCR, but not the P1, might be sensitive to the manipulation of stimulus duration, we added the fixed-effect *duration* (100 ms versus 1000 ms) in the SCR LMM.

#### Exploratory analyses

To further explore the retrieval phase, we accounted for contingency awareness. Previously, contingency awareness has been shown to mediate threat responses in the SCR (Sevenster et al., 2014; Tanskanen et al., 2007). We also conducted these analyses for the P1 data, although we did not have any *a priori* assumptions regarding the analysis outcome. To avoid overspecifying the model, we analyzed the models reported in the paragraph above for each group (contingency aware/contingency unaware) separately. Additionally, we compared overall P1 and SCR amplitudes in the acquisition and retrieval phase between contingency aware and unaware participants with non-hierarchical linear regression models. The results are reported in the supplementary analyses of P1 and SCR data, respectively.

#### EEG-SCR association

To test for the association of P1 and SCR data, we calculated an additional LMM, predicting trial-wise SCR values based on P1 amplitudes in interaction with the fixed-effects *CS*, *SF*, and *hemisphere*. Before analysis, both P1 and SCR data were *z*-transformed. For post-hoc tests of regression slopes, unstandardized beta coefficients are reported as effect sizes in addition to the Std-β. Only trials that were present both in the P1 and SCR analysis were considered for analysis, and the number of trials was balanced between *CS* conditions (see above). The LMM was specified with the additive fixed-effect *trial* to avoid inflating the correlation of SCR and P1 amplitudes by potentially simultaneous signal habituation (Lonsdorf et al., 2017).

### Data availability

All data needed to evaluate the conclusions are present in the paper or the supporting information. The datasets generated in this study are available under this link: https://gitlab.ub.uni-bielefeld.de/ae02/weidner_subcortical_threat_retrieval so are numerical source data for graphs and charts. Raw data are available from the authors upon reasonable request to the corresponding author.

### Code availability

The codes generated in this study are available under this link: https://gitlab.ub.uni-bielefeld.de/ae02/weidner_subcortical_threat_retrieval.

## Results

### Stimulus ratings

Both rating scales differentiated threat from safety, with lower valence (β = -0.804, *p* = 7.12e^-06^. Power (*M*[*CI*]) = 99.60 (98.56, 99.95) %) and higher arousal scores for CS+ than CS- (β = 0.449, *p* = .004, Power (*M*[*CI*]) = 85.40 (82.00, 88.38) %). Separating the analyses based on contingency awareness showed that the CS-differentiation in both rating scales was only present for contingency aware participants (arousal: β = 0.956, *p* = 1.18e^-05^; valence: β = -1.456, *p* = 4.31e^-08^) while contingency unaware participants did not show differential ratings based on the *CS* condition (arousal: β = -0.043, *p* = .831; valence: β = -0.171, *p* = .463). Figure 1B illustrates the rating results of all participants. Full model coefficients are found in Table S1.

### P1

#### Acquisition: Broadband presentation

The LMM indicated a significant fixed-effect *hemisphere* during the acquisition phase, with higher P1 amplitudes over the left than the right hemisphere (β = -0.314, *p* = .028; Power (*M*[*CI*]) = 78.40 (74.53, 81.93) %). No significant effect of *CS* was observed (β = -0.122, *p* = .395). The *CS* × *hemisphere* interaction was also not significant (β = 0.024, *p* = .905). Full model coefficients are found in Table S2. Separate P1 analyses with the subsample of participants who also had available SCR data confirmed the same pattern of activity as found in the full sample (see Table S6). The pattern of results did not change when separating the analyses based on contingency awareness (see Table S2), although the effect of hemisphere was not present in contingency unaware participants (β = 0.003, *p* = .990).

#### Retrieval: Filtered presentation

The LMM revealed significant differences in the P1 amplitude during the retrieval phase between CS+ and CS- (CS+ > CS-; β = 0.205, *p* = .029, Power (*M*[*CI*]) = 86.40 (83.08, 89.28) %, Fig. 2). Furthermore, the three-way interaction *CS* × *SF* × *hemisphere* was significant (β = 0.409, *p* = .029, Power (*M*[*CI*]) = 60.20 (55.76, 64.52) %). Post-hoc tests revealed that this was driven by lateralized changes in CS-differentiation based on spatial frequency: The largest CS-differentiation was found for LSF faces over the left hemisphere (β = -0.268, *p* = .012, *d* = -0.326). Over the right hemisphere, marginally larger P1 amplitudes for CS+ versus CS- faces were found for HSF faces (β = -0.205, *p* = .058, *d* = -0.244). No significant CS-differences were observed for LSF over the right or for HSF over the left hemisphere (Fig. 3). Full model coefficients are found in Table 1, post-hoc tests are found in Table S3. Separate P1 analyses with the subsample of participants who also had available SCR data confirmed the same pattern of activity as found in the full sample (see Table S7 and Table S8).

**Figure 2.**
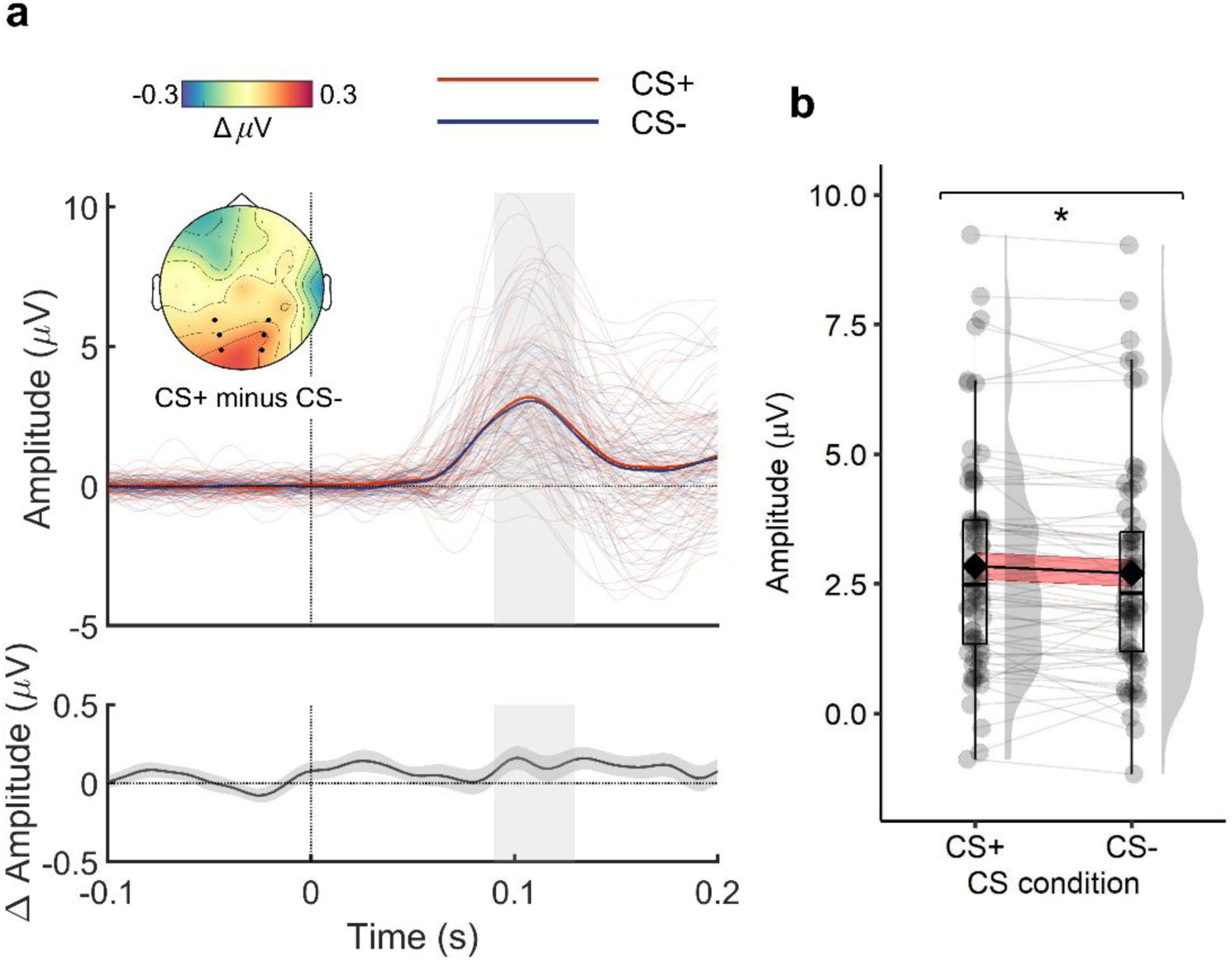
CS differentiation in P1 amplitude. **a** Average evoked time-series and differential scalp topography of responses during the P1-time window (90 – 130 ms). Channels that were used to extract the P1 are marked in black. Thin lines depict single-subject time-series per *CS* condition. The shaded area shows the P1 time-window. **b** Mean ERP amplitudes in response to the *CS* conditions. Distribution data are averaged across P1 time-points and channels. Boxplots show median and interquartile range; raincloud plots display single-subject means per *CS* condition along with the smoothed distribution density; diamonds indicate the overall condition mean. Line plots depict data trends. The black line shows the mean across all subjects; red ribbons indicate standard error of the mean; grey lines show single-subject trends. Stars and brackets mark significant post-hoc tests (\**p* ≤ .05). *Abbreviations*: CS = conditioned stimulus.

**Figure 3.**
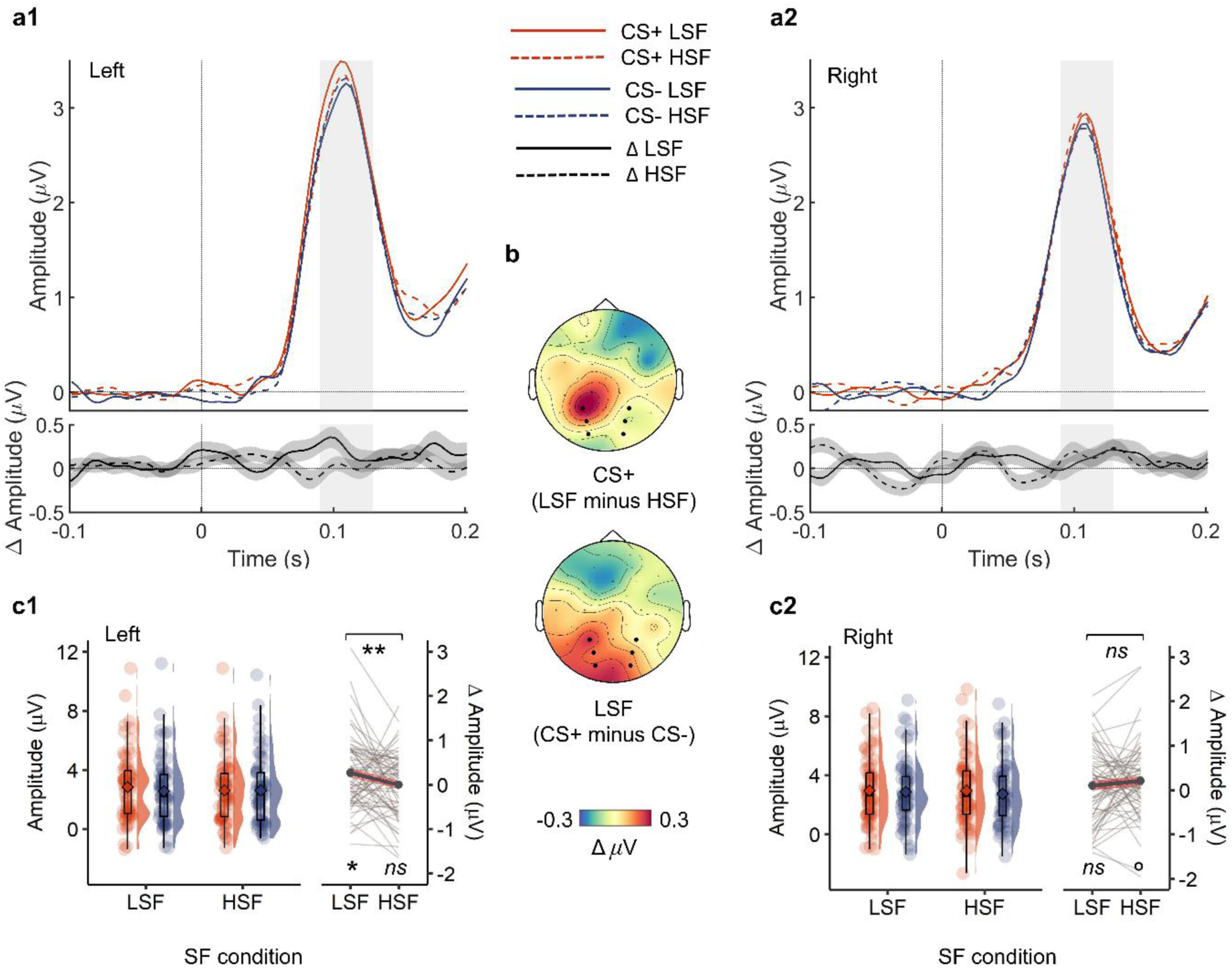
CS-differentiation in P1 amplitude as a function of spatial frequency and hemisphere. **a** Average and differential (CS+ versus CS-) evoked time-series of responses in the left (a1) and right (a2) channel cluster. The shaded area marks the P1 time-window. **b** Differential (CS+ versus CS-) scalp topography during the P1-time window (90 – 130 ms). Channels that were used to extract the P1 are marked in black. **c** Distribution of P1 amplitudes in response to each *CS* × *SF* condition for the left (c1) and right (c2) hemisphere. Boxplots show median and interquartile range; raincloud plots display single-subject means per *CS* × *SF* condition along with the smoothed distribution density; outlined diamonds indicate the overall condition mean. Line plots depict the CS-differentiation (CS+ minus CS-) within *SF* conditions. Bold grey lines indicate mean group differences; red ribbons indicate standard error of the difference; light grey lines depict single-subject mean differences. Stars and brackets mark significant post-hoc tests (°*p* ≤ .10, \**p* ≤ .05, \*\**p* ≤ .01). *Abbreviations*: CS = conditioned stimulus, LSF = low spatial frequency, HSF = high spatial frequency, ns = not significant, SF = spatial frequency.

**Table 1.** P1 – Retrieval: Linear Mixed Model Fixed Effects.

| Term | Estimate | SE | df | t | p |
| --- | --- | --- | --- | --- | --- |
| *Full sample* |  |  |  |  |  |
| (Intercept) | **2.743** | **0.257** | **75.482** | **10.665** | **1.01e<sup>-16</sup>** |
| CS | **0.205** | **0.093** | **47,005.211** | **2.189** | **.029** |
| SF | 0.145 | 0.094 | 47,005.393 | 1.554 | .120 |
| Hemisphere | -0.121 | 0.093 | 47,005.269 | -1.296 | .195 |
| CS×SF | -0.131 | 0.132 | 47,005.325 | -0.993 | .321 |
| CS×Hemisphere | -0.214 | 0.132 | 47,005.138 | -1.620 | .105 |
| SF×Hemisphere | -0.201 | 0.132 | 47,005.271 | -1.517 | .129 |
| CS×SF×Hemisphere | **0.409** | **0.187** | **47,005.139** | **2.185** | **.029** |
| *Contingency aware* |  |  |  |  |  |
| (Intercept) | **3.461** | **0.390** | **35.168** | **8.873** | **1.69e<sup>-10</sup>** |
| CS | 0.131 | 0.127 | 23,136.057 | 1.031 | .288 |
| SF | 0.097 | 0.127 | 23,136.108 | 0.763 | .451 |
| Hemisphere | 0.132 | 0.127 | 23,136.102 | 1.042 | .302 |
| CS×SF | -0.137 | 0.180 | 23,136.085 | -0.762 | .434 |
| CS×Hemisphere | -0.306 | 0.180 | 23,136.049 | -1.700 | .089 |
| SF×Hemisphere | -0.315 | 0.180 | 23,136.078 | -1.756 | .081 |
| CS×SF×Hemisphere | **0.647** | **0.254** | **23,136.037** | **2.549** | **.013** |
| *Contingency unaware* |  |  |  |  |  |
| (Intercept) | **2.068** | **0.295** | **41.524** | **7.011** | **1.49e<sup>-08</sup>** |
| CS | **0.273** | **0.137** | **23,862.230** | **2.000** | **.044** |
| SF | 0.192 | 0.137 | 23,862.447 | 1.400 | .154 |
| Hemisphere | **-0.366** | **0.137** | **23,862.251** | **-2.674** | **.009** |
| CS×SF | -0.119 | 0.194 | 23,862.343 | -0.613 | .542 |
| CS×Hemisphere | -0.122 | 0.193 | 23,862.144 | -0.633 | .523 |
| SF×Hemisphere | -0.090 | 0.194 | 23,862.304 | -0.464 | .652 |
| CS×SF×Hemisphere | 0.167 | 0.274 | 23,862.170 | 0.610 | .546 |

*Note.* *T*-statistics are evaluated for significance. A positive coefficient indicates higher ratings in the test condition than in the reference condition. Effects with $p \leq .05$ are marked in bold. *Abbreviations:* CS = conditioned stimulus, SE = standard error, SF = spatial frequency.

When separating the analyses based on contingency awareness, the *CS* × *SF* × *hemisphere* interaction was only present for contingency aware participants (contingency aware: β = 0.647, *p* = .013; contingency-unaware: β = 0.167, *p* = .546). P1 amplitudes from contingency unaware participants showed a significant effect of *CS* (CS+ > CS-; β = 0.273, *p* = .044) as well as a significant effect of *hemisphere* (right > left; β = -0.366, *p* = .009) but no difference depending on *SF*. Full model coefficients are found in Table 1, post-hoc tests are found in Table S3.

### SCR

#### Acquisition: Broadband presentation

Investigating the SCR during the acquisition phase, the LMM revealed, just as for the P1, no significant effect of *CS* (β = 0.00003, *p* = .930). Full model coefficients are found in Table S4. The pattern of results did not change when separating the analyses based on contingency awareness (see Table S4).

#### Retrieval: Filtered presentation

As shown in Fig. 4a, the LMM revealed a significant *CS* × *duration* interaction (β = 0.017, *p* = .021; Power (*M*[*CI*]) = 46.60 (42.16, 51.08) %). CS-differentiation increased for longer (1000 ms) stimulus durations, although this reached only marginal significance in the Holm-corrected post-hoc tests (1000 ms: CS- versus CS+: β = -0.007, *p* = .071, *d* = -0.179; 100 ms: CS- versus CS+: β = 0.005, *p* = .218, *d* = 0.259). The model also revealed a significant effect of *duration* (100 ms > 1000 ms; β = -0.012, *p* = .022; Power (*M*[*CI*]) = 48.80 (44.34, 53.28) %) as well as a significant *SF* × *duration* interaction (β = 0.016, *p* = .035; Power (*M*[*CI*]) = 51.60 (47.12, 56.60) %). Post-hoc tests showed that SCR amplitudes marginally increased in response to LSF HSF faces when shown for 1000 ms (1000 ms: LSF > HSF: β = -0.007, *p* = .072, *d* = -0.255; 100 ms: LSF = HSF: β = 0.003, *p* = .342, *d* = 0.182). Full model coefficients are found in Table 2, post-hoc tests are found in Table S5.

**Figure 4.**
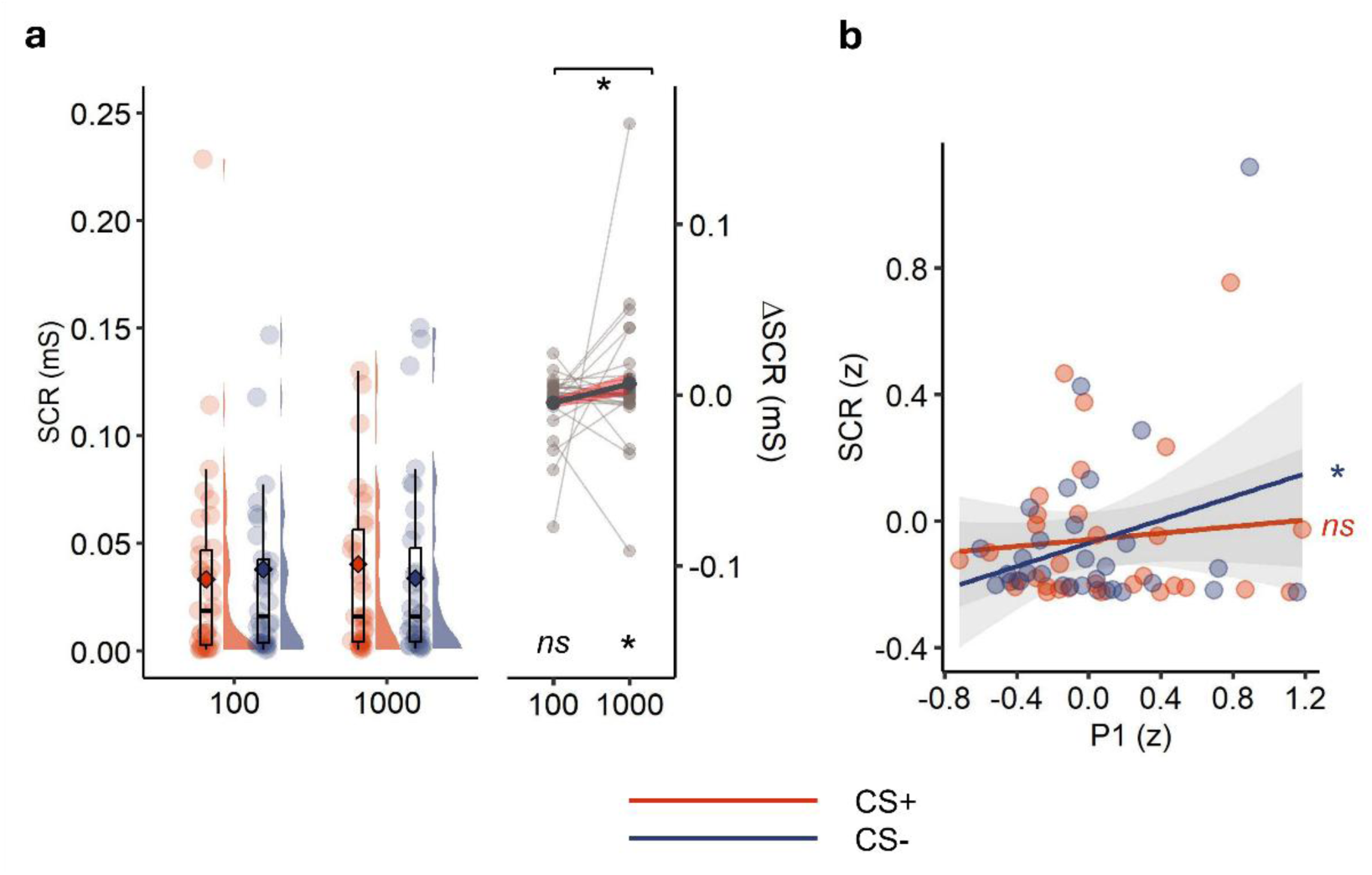
CS-differentiation in SCR and association with P1 amplitudes. **a** SCR amplitude in response to the *CS* × *duration* conditions. Boxplots show median and interquartile range; raincloud plots display single-subject means per *CS* × *duration* condition along with the smoothed distribution density; diamonds indicate the overall condition mean. Line plots depict the CS-differentiation (CS+ versus CS-) within duration conditions. The bold grey line indicates mean group differences; red ribbons indicate standard error of the difference; light grey lines depict single-subject mean differences. **b** *Z*-transformed P1-SCR associations within the *CS* conditions. Line plots depict standardized linear regression slopes of P1 amplitude based on SCR peak amplitude along with the standard error (shaded area). Dots display single-subject means per *CS* condition. Stars and brackets mark significant post-hoc tests (\**p* ≤ .05). *Abbreviations*: CS = conditioned stimulus, ns = not significant, SCR = skin conductance response.

**Table 2.** SCR – Retrieval: Linear Mixed Model Fixed Effects.

| Term | Estimate | SE | df | t | p |
| --- | --- | --- | --- | --- | --- |
| *Full sample* |  |  |  |  |  |
| (Intercept) | **0.042** | **0.009** | **44.23** | **4.576** | **.002** |
| CS | -0.010 | 0.005 | 12,125.80 | -1.847 | .069 |
| SF | -0.009 | 0.005 | 12,125.60 | -1.672 | .090 |
| Duration | **-0.012** | **0.005** | **12,125.86** | **-2.282** | **.022** |
| CS×SF | 0.010 | 0.007 | 12,125.78 | 1.388 | .167 |
| CS×Duration | **0.017** | **0.007** | **12,126.38** | **2.256** | **.021** |
| SF×Duration | **0.016** | **0.007** | **12,126.24** | **2.118** | **.035** |
| CS×SF×Duration | -0.011 | 0.010 | 12,126.21 | -1.053 | .303 |
| *Contingency aware* |  |  |  |  |  |
| (Intercept) | **0.046** | **0.014** | **25.216** | **3.428** | **.002** |
| CS | -0.009 | 0.007 | 7,552.344 | -1.323 | .198 |
| SF | -0.008 | 0.007 | 7,552.295 | -1.227 | .210 |
| Duration | **-0.019** | **0.007** | **7,552.312** | **-2.795** | **.003** |
| CS×SF | 0.011 | 0.010 | 7,552.310 | 1.161 | .254 |
| CS×Duration | **0.025** | **0.010** | **7,552.627** | **2.562** | **.011** |
| SF×Duration | **0.020** | **0.010** | **7,552.689** | **2.050** | **.035** |
| CS×SF×Duration | -0.015 | 0.014 | 7,552.603 | -1.113 | .271 |
| *Contingency unaware* |  |  |  |  |  |
| (Intercept) | **0.036** | **0.011** | **20.707** | **3.351** | **.003** |
| CS | -0.01 | 0.008 | 4,566.642 | -1.326 | .178 |
| SF | -0.009 | 0.008 | 4,566.418 | -1.167 | .226 |
| Duration | -9.87e<sup>-05</sup> | 0.008 | 4,566.811 | -0.013 | .992 |
| CS×SF | 0.008 | 0.011 | 4,566.655 | 0.739 | .478 |
| CS×Duration | 0.003 | 0.011 | 4,567.053 | 0.280 | .776 |
| SF×Duration | 0.009 | 0.011 | 4,566.657 | 0.801 | .418 |
| CS×SF×Duration | -0.004 | 0.016 | 4,566.736 | -0.252 | .825 |

Note. *T*-statistics are evaluated for significance. A positive coefficient indicates higher ratings in the test condition than in the reference condition. Effects with $p \leq .05$ are marked in bold. Abbreviations: CS = conditioned stimulus, SCR = skin conductance response, SE = standard error, SF = spatial frequency.

When separating the analyses based on contingency awareness, the abovementioned *CS* × *duration* interaction in the SCR was only, and more strongly, present for contingency aware participants (contingency aware: β = 0.025, *p* = .011; contingency unaware: β = 0.003, *p* = .776). Here, the post-hoc comparison of CS- versus CS+ reached significance when stimuli were presented for 1000 ms (CS+ > CS-, β = -0.014, *p* = .008. *d* = -0.367). Likewise, the *SF* × *duration* was only found for contingency aware participants (contingency aware: β = 0.020, *p* = .035; contingency unaware: β = 0.009, *p* = .418). For those participants, SCR amplitude increased more strongly in response to LSF than HSF faces only when shown for 1000 ms (1000 ms: LSF > HSF: β = -0.010, *p* = .49, *d* = -0.354; 100 ms: LSF = HSF: β = 0.003, *p* = .579, *d* = 0.148). Full model coefficients are found in Table 2, post-hoc tests are found in Table S5.

### EEG-SCR association

The model revealed a significant prediction of SCR amplitudes by P1 amplitudes, even after accounting for trial-wise habituation (EEG: β = 0.195, *p* = .035, Fig. 4a). As shown in Figure 4b, this effect interacted marginally with the CS condition (*EEG* × *CS*: β = -0.212, *p* = .077). Post-hoc tests revealed that the only significant association of SCR amplitudes with P1 amplitudes was present for CS- faces (CS-: Std-β = .033, β = 0.001, *p* = .007; CS+: Std-β = .013, β = 0.0006, *p* = .285). Full model coefficients are found in Table S9, post-hoc tests are found in Table S10.

Separating the analyses based on contingency awareness, we found that the abovementioned association of P1 and SCR amplitudes as well as the *EEG* × *CS* interaction was only present for contingency-unaware participants (unaware: EEG: β = 0.171, *p* = .050; *EEG* × *CS*: β = -0.222, *p* = .040; aware: EEG: β = 0.212, *p* = .070; *EEG* × *CS*: β = -0.207, *p* = .205). However, post-hoc tests showed that neither a P1-SCR association for CS- nor for CS+ faces was significant in contingency unaware participants (CS-: Std-β = .024, β = 0.001, *p* = .237; CS+: Std-β = -.008, β = -0.0006, *p* = .693). Full model coefficients are found in Table S9, post-hoc tests are found in Table S10.

## Discussion

Previous studies suggest that detection of facial threat in early visual attention, as reflected in the P1 component, may be driven by LSF visual information. However, it remained unclear whether any such effects, often investigated by comparing responses to fearful versus neutral faces, are due to configural differences between expressions or genuine detection of threat relevance (Bruchmann et al., 2020; Schindler, Bruchmann, et al., 2021; Schindler, Wolf, et al., 2021; Yu et al., 2023). Likewise, it was unknown whether SF-specific effects of fear retrieval are present for novel threat associations. To address these issues, we investigated the contribution of SF characteristics to threat detection in fear-conditioned faces with neutral expressions. Furthermore, we explored whether SF-specific P1 modulations are reflected in autonomic arousal.

First, we could confirm that threat detection in the P1 component is retained even in the absence of inherent physical cues of threat-specific facial expressions. This contradicts previous hypotheses suggesting that mainly perceptual factors underlie P1 sensitivity to threat-related faces, although they might play a modulatory role (Schindler, Wolf, et al., 2021). Indeed, confirming our pre-registered hypotheses, P1 threat retrieval was distinctly shaped by the spatial frequency content of the face, albeit in a lateralized manner. P1 amplitudes showed largest amplitudes in response to LSF CS+ faces over the left hemisphere, whereas over the right hemisphere, threat detection was, in tendency, mediated by HSF. Interestingly, although general CS-differentiation occurred irrespective of contingency awareness, LSF-specific CS-differentiation was only present in contingency aware participants. Against our hypotheses, SCR sensitivity towards CS+ faces was not driven by SF but by long stimulus durations and contingency awareness. In contingency unaware participants, P1 and SCR conditioning effects were positively correlated. CS-differentiation during the acquisition phase was absent in both the P1 and SCR, potentially because too few trials with established US-contingencies were available (Bach et al., 2023).

P1 data from the retrieval phase support the assumption that magnocellular signals can drive the retrieval of threat associations at the earliest stages of visuocortical processing (Carretié et al., 2017; Méndez-Bértolo et al., 2016; Morris et al., 1999), even when faces are perceptually neutral. They highlight the ability of magnocellular-cortical pathways to adapt remarkably quickly (Cinca-Tomás et al., 2025; Weinberger, 1982; Zhuang et al., 2015) and suggest that magnocellular learning might underlie the initial formation of threat associations. Notably, CS-differentiation of LSF faces effect was constrained to the left hemisphere. This contrasts the well-known right-hemispheric dominance in face processing (Haxby et al., 2000; Kanwisher et al., 1997; Kanwisher & Yovel, 2006), and suggests additional left lateralized mechanisms of threat perception that are specific to LSF information. Indeed, particularly visual discrimination in the left hemisphere has been linked to LSF processing (Kitterle et al., 1990; Peyrin et al., 2003) and magnocellular signaling (Okubo & Nicholls, 2005). More specifically, the presently observed temporo-parietal focus of SF differences for CS+ faces could point towards left-dominant magnocellular projections to the dorsal visual stream (Bullier, 2001; Maunsell et al., 1990; Nassi & Callaway, 2006; Skottun, 2015; Tootell & Nasr, 2017) potentially informing prediction error coding in the parietal cortex (Spoormaker et al., 2011). In line with this, Steinberg et al. (2012) observed differentiation of fear conditioned faces at 130 – 190 ms to spread from occipito-temporal to left parietal cortices. Following Carretié et al. (2020), it is likely that threat information is directly evaluated within magnocellular subdivisions projecting to the dorsal visual stream and entrained by the amygdala (Cinca-Tomás et al., 2025; Maren et al., 2001). As the dorsal visual stream is implicated in attention and spatial orientation (Corbetta et al., 2008; Goodale & Milner, 1992; Milner & Goodale, 2006), its connections to the magnocellular divisions of the thalamus might support rapid, attentional engagement with the fear-associated stimulus (Pourtois et al., 2005). Indeed, saccadic activity was observed to be faster for threatening than neutral images when preceded by a LSF prime (Zhu et al., 2021). The present data further suggest that CS-differentiation of LSF faces is mediated by contingency awareness. Similarly, during auditory processing, Kosteletou Kassotaki et al. (in press) recently reported an association between the density of magnocellular-amygdala fiber tracts and self-reported fearfulness. Potentially, magnocellular threat processing promotes contingency learning because it quickly engages attention (Bullier, 2001; Maunsell et al., 1990; Nassi & Callaway, 2006; Skottun, 2015; Tootell & Nasr, 2017), as also implied in higher P1 amplitudes in contingency aware than unaware participants. In turn, contingency awareness might also contribute to perceptual learning (Kuchinke et al., 2015): The frontal cortex that potentially mediates contingency awareness (Baeuchl et al., 2019; Kuchinke et al., 2015; Madaboosi et al., 2021; Tabbert et al., 2011) could have also guided associative learning in magnocellular pathways (Mechias et al., 2010; Mitchell et al., 2007) and CS-differentiation in the P1 (Kuchinke et al., 2015). Of note, in the present study, CS-differentiation was still found in contingency unaware participants albeit not specific to SF characteristics. Therefore, some P1 CS-differentiation can occur without conscious contingency awareness. A driving node for this could be the amygdala, which can acquire US-contingencies without conscious awareness (Tabbert et al., 2011). Correspondingly, amygdala activity in fMRI studies was found positively associated with P1 amplitudes (Müller-Bardorff et al., 2018).

In tendency, right-hemispheric P1 amplitudes showed CS-differentiation for HSF faces. As a speculative explanation, this trend may imply a stronger reliance on parvocellular threat-signals in the right hemisphere. It could also reflect the acquisition of threat-associations in right-hemispheric lateral occipital regions (Li & Keil, 2023; Steinberg et al., 2012). As part of the right-dominant face processing network (Fox et al., 2009; Kanwisher et al., 1997; Kanwisher & Yovel, 2006), these regions are implicated in structural encoding of faces (Cesarei et al., 2013; Schindler, Bruchmann, et al., 2021) and may rely on detailed HSF information (Rebaï et al., 1998; Zhao et al., 2023). Therefore, it appears that multiple, parallel pathways of threat processing adapt to distinct visual features during learning (Bruchmann et al., 2023; Denison & Silver, 2012).

In the SCR, heightened autonomic arousal in response to CS+ than CS- faces was observed only for faces that were presented for longer durations. Increased time to engage with the stimulus might have improved associative learning because it allows for more detailed categorization of facial identity (Tanskanen et al., 2007) which was the key-predictor of US-contingency in the present study. Correspondingly, greater SCR sensitivity to CS+ than CS- was linked to associative processing in frontal areas (Andreatta et al., 2015; Carter et al., 2006; McIntosh et al., 1999; Mechias et al., 2010). In parallel, longer presentation times were also associated with higher SCR to LSF than HSF faces. Park and colleagues (2016) argue that, due to the assumed involvement of the amygdala in the magnocellular pathway, particularly LSF stimuli elicit higher arousal. Correspondingly, the categorization of LSF identities was previously shown to be skewed towards negative interpretations (Neta & Whalen, 2010; Park et al., 2016). In conjunction with the observation that contingency-awareness drove the CS-differentiation, the present findings suggest that the SCR reflects conscious anticipatory arousal. Indeed, previous work has demonstrated that threat differentiation in the SCR emerged most robustly when participants were aware of the US-contingencies (Baeuchl et al., 2019; Sevenster et al., 2014). Interestingly, particularly in contingency unaware participants, physiological arousal appeared to track P1 amplitudes. Hence, attentional orienting in the P1 seemingly informed arousal more than in contingency aware participants, potentially due to inefficient associative learning (Baeuchl et al., 2019; Kuchinke et al., 2015; Madaboosi et al., 2021; Tabbert et al., 2011). Given that the P1-SCR association was additionally more positive for CS- than CS+ faces in unaware participants, inefficient learning might underlie threat-overgeneralization. Similarly, responses to CS- are often more predictive of anxiety disorders than responses to CS+ as these patients present defective safety learning (Lissek, 2012).

Despite these novel insights into threat retrieval, some limitations must be considered when interpreting the present findings. First, the localization of the CS-differentiation remains imprecise due to the sparse electrode coverage on the scalp, which limits conclusions about the underlying neural generators. While we associated effects of LSF CS+ with parietal regions of the dorsal visual stream, contributions from frontal areas could also be present (e.g., Rehbein et al., 2014). Frontal regions are implicated in contingency learning (Mechias et al., 2010) and magnocellular–frontal connections have been linked to the acquisition of stimulus-stimulus associations (Mitchell et al., 2007). Future studies employing higher-density recordings and source localization could address the underlying spatio-temporal mechanisms. Furthermore, it is unclear how SF-specific threat differentiation would evolve beyond the present testing interval. For example, threat associations could be increasingly represented in HSF information through cortical learning (Li & Keil, 2023; McTeague et al., 2015). Potentially, there is first evidence for this mechanism in the present right-hemispheric trend for CS-differentiation in HSF. Additionally, effects of contingency awareness were explored without direct experimental modulation. Future studies should revisit this and test the replicability of effects.

Taken together, this study provides compelling evidence that the magnocellular facilitation of visuocortical threat detection in faces goes beyond pre-existing configural clues of specific facial expressions. It also indicates the presence of several parallel processes that differentially modulate early visual processing and autonomic arousal in response to potential threat, although both P1 and SCR effects shared their association with contingency awareness. These findings refine models of emotional vision and demonstrate that the P1 component can reflect rapid acquisition of threat associations in a SF-specific manner. Understanding the neural circuits that underlie fear retrieval holds important implications for research on maladaptive fear responses found in trauma-related (Careaga et al., 2016) or anxiety disorders (Chien et al., 2020; Kosteletou Kassotaki et al., in press). Research should consider the different neural circuits of visual processing and their timing during fear retrieval. In doing so, it could potentially inform therapeutic protocols, such as neuromodulation techniques (Dittert et al., 2018; Zhang et al., 2024), exposure therapy (Lange et al., 2020; Winkler et al., 2025), or neurofeedback (Nicholson et al., 2017).

## Acknowledgements

Thank you to Mana Ehlers for providing a preprocessing and analysis framework for the SCR data. Thank you to Sebastian Geukes for helping with the LMM pipeline. We would like to express our gratitude to all the participants.

## Ethics approval

The study was conducted in accordance with the Declaration of Helsinki and was approved by the Ethics committee of Bielefeld University (EUB-2024_245), following the ethics guidelines of the German Psychological Association (DGPs).

## Competing Interest Statement

The authors declare no competing financial interests.

## CrediT statement

EW: conceptualization, project administration, data curation, formal analysis, methodology, visualization, writing – original draft, writing – review & editing; AT and MG: investigation, data curation, writing – review & editing; JK: supervision, resources, writing – review & editing

## Supplementary Information

### SAM ratings

**Table S1.** SAM Ratings – Linear Mixed Model Fixed Effects.

| Scale | Term | Estimate | SE | df | t | p |
| --- | --- | --- | --- | --- | --- | --- |
| Arousal | *Full sample* |  |  |  |  |  |
|  | (Intercept) | **4.297** | **0.135** | **171.010** | **31.778** | **1.16e<sup>-73</sup>** |
|  | CS | **-0.804** | **0.175** | **206.00** | **-4.608** | **7.12e<sup>-06</sup>** |
|  | *Contingency aware* |  |  |  |  |  |
|  | (Intercept) | **2.647** | **0.213** | **54.839** | **12.404** | **1.51e<sup>-17</sup>** |
|  | CS | **0.956** | **0.207** | **101.000** | **4.609** | **1.18e<sup>-05</sup>** |
|  | *Contingency unaware* |  |  |  |  |  |
|  | (Intercept) | **2.943** | **0.204** | **57.075** | **14.429** | **1.08e<sup>-20</sup>** |
|  | CS | -0.043 | 0.200 | 104.000 | -0.214 | .831 |
| Valence | *Full sample* |  |  |  |  |  |
|  | (Intercept) | **2.797** | **0.148** | **116.319** | **18.881** | **3.21e<sup>-37</sup>** |
|  | CS | **0.449** | **0.148** | **206.000** | **3.039** | **.004** |
|  | *Contingency aware* |  |  |  |  |  |
|  | (Intercept) | **4.794** | **0.178** | **94.24** | **26.872** | **5.73e<sup>-46</sup>** |
|  | CS | **-1.456** | **0.246** | **101.00** | **-5.927** | **4.31e<sup>-08</sup>** |
|  | *Contingency unaware* |  |  |  |  |  |
|  | (Intercept) | **3.814** | **0.195** | **74.398** | **19.520** | **5.39e<sup>-31</sup>** |
|  | CS | -0.171 | 0.233 | 104.000 | -0.736 | .463 |

Notes. Ratings were given on a 7-point Likert scale, 1 being most negative/least arousing. *T*-statistics are evaluated for significance. A positive coefficient indicates higher ratings in the test condition than in the reference condition. Effects with $p \leq .05$ are marked in bold. Abbreviations: CS = conditioned stimulus, SE = standard error.

### P1 analysis

#### Contingency awareness: General amplitude differences

The non-hierarchical linear models revealed higher P1 amplitudes for contingency aware than unaware participants in the acquisiton phase (β = 1.245, *t*(66), *p* = .019) as well as in the retrieval phase (β = 1.478, *t*(66) = 3.149, *p* = .002).

**Table S2.**
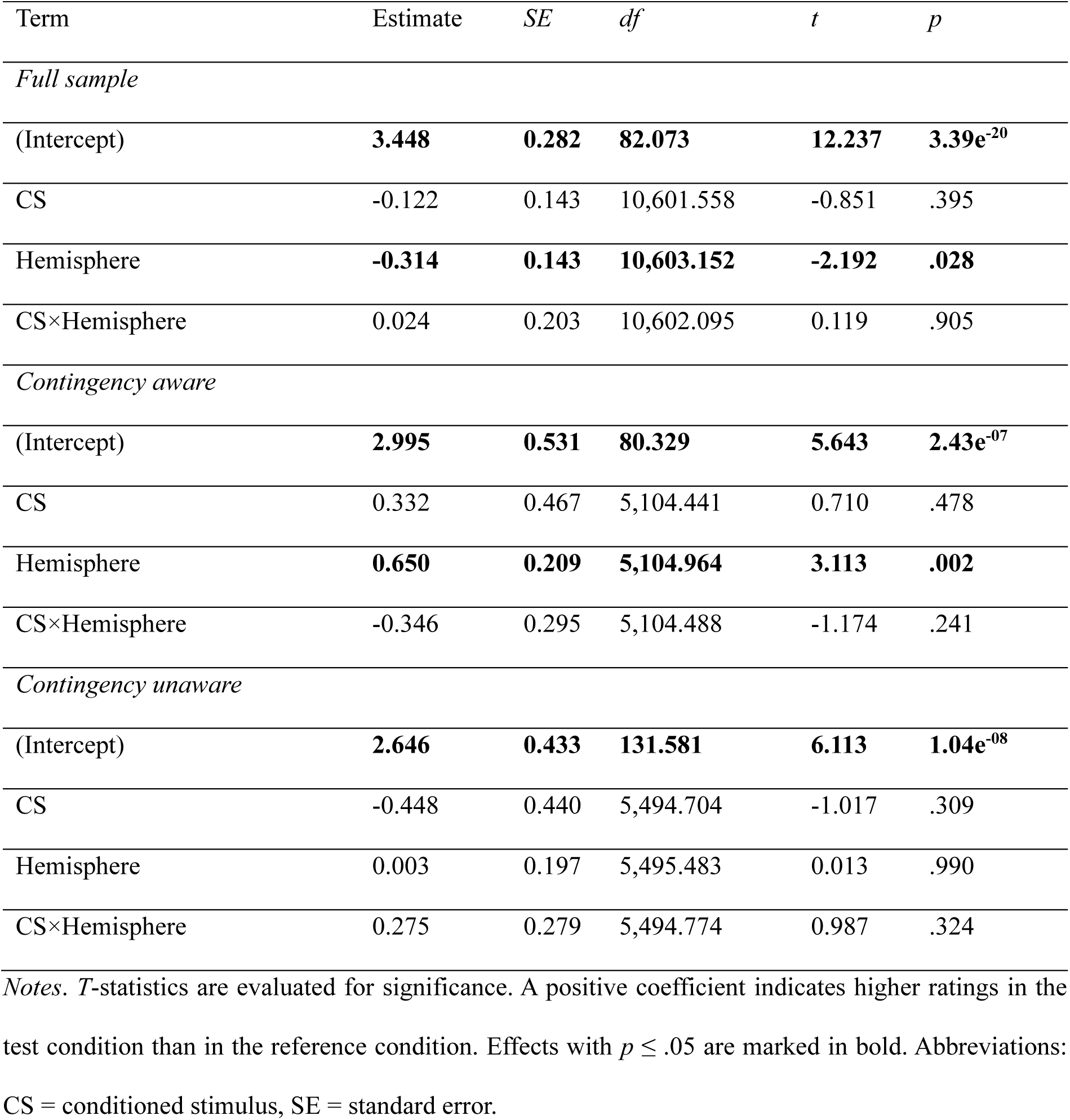
P1 Acquisition: Linear Mixed Model Fixed Effects.

**Table S3.** P1 Retrieval: Post-hoc CS Differences by SF × Hemisphere.

| SF | Hemisphere | Contrast | Estimate | SE | df | z | p | d |
| --- | --- | --- | --- | --- | --- | --- | --- | --- |
| *Full sample* |  |  |  |  |  |  |  |  |
| HSF | Left | (CS-) - (CS+) | 0.009 | 0.093 | 47080 | 0.100 | .920 | 0.004 |
| LSF | Left | (CS-) - (CS+) | **-0.268** | **0.094** | **47080** | **-2.862** | **.012** | **-0.326** |
| HSF | Right | (CS-) - (CS+) | -0.205 | 0.093 | 47080 | -2.189 | .058 | -0.244 |
| LSF | Right | (CS-) - (CS+) | -0.073 | 0.094 | 47080 | -0.784 | .433 | -0.116 |
| *Contingency aware* |  |  |  |  |  |  |  |  |
| HSF | Left | (CS-) - (CS+) | 0.175 | 0.127 | 23904 | 1.373 | .170 | 0.293 |
| LSF | Left | (CS-) - (CS+) | **-0.336** | **0.127** | **23904** | **-2.649** | **.008** | **-0.421** |
| HSF | Right | (CS-) - (CS+) | -0.131 | 0.127 | 23904 | -1.031 | .303 | -0.181 |
| LSF | Right | (CS-) - (CS+) | 0.006 | 0.127 | 23904 | 0.046 | .964 | 0.010 |

*Notes.* *T*-statistics are evaluated for significance. A positive coefficient indicates higher ratings in the test condition than in the reference condition. Effects with $p \leq .05$ are marked in bold. Abbreviations: CS = conditioned stimulus, HSF = high spatial frequency, LSF = low spatial frequency, SE = standard error, SF = spatial frequency.

### SCR

#### Contingency awareness: General amplitude differences

The non-hierarchical linear models revealed no differences in SCR amplitudes betweem contingency aware and unaware participants in the acquisiton phase (β = 0.024, *t*(32), *p* = .194) as well as in the retrieval phase (β = -0.010, *t*(32) = -0.558, *p* = .581).

**Table S4.** SCR Acquisition: Linear Mixed Model Fixed Effects.

| Term | Estimate | SE | df | t | p |
| --- | --- | --- | --- | --- | --- |
| *Full sample* |  |  |  |  |  |
| (Intercept) | 0.021 | 0.167 | 2,909.437 | 0.125 | .901 |
| CS | $3.03\text{e}^{-04}$ | 0.003 | 2,900.020 | 0.088 | .930 |
| *Contingency aware* |  |  |  |  |  |
| (Intercept) | **0.049** | **0.013** | **26.154** | **3.846** | **$6.93\text{e}^{-04}$** |
| CS | 0.004 | 0.006 | 1,879.018 | 0.660 | .509 |
| *Contingency unaware* |  |  |  |  |  |
| (Intercept) | **0.031** | **0.010** | **14.875** | **3.116** | **.007** |
| CS | -0.009 | 0.006 | 1,020.009 | -1.410 | .159 |

*Notes.* T-statistics are evaluated for significance. A positive coefficient indicates higher ratings in the test condition than in the reference condition. Abbreviations: CS = conditioned stimulus, SE = standard error.

**Table S5.**
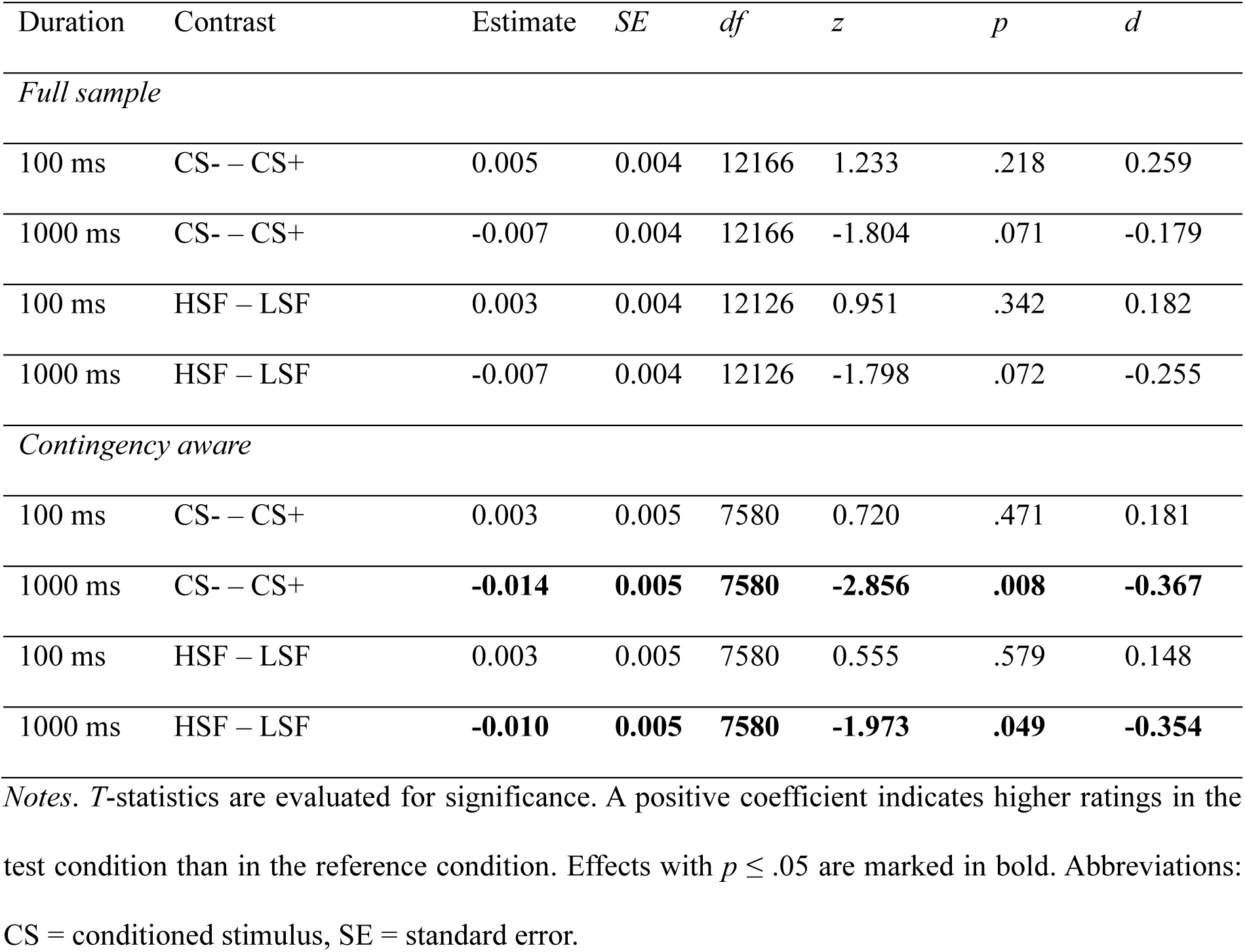
SCR Retrieval: Post-hoc CS/SF Differences by Duration.

| Duration | Contrast | Estimate | SE | df | z | p | d |
| --- | --- | --- | --- | --- | --- | --- | --- |
| *Full sample* |  |  |  |  |  |  |  |
| 100 ms | CS- – CS+ | 0.005 | 0.004 | 12166 | 1.233 | .218 | 0.259 |
| 1000 ms | CS- – CS+ | -0.007 | 0.004 | 12166 | -1.804 | .071 | -0.179 |
| 100 ms | HSF – LSF | 0.003 | 0.004 | 12126 | 0.951 | .342 | 0.182 |
| 1000 ms | HSF – LSF | -0.007 | 0.004 | 12126 | -1.798 | .072 | -0.255 |
| *Contingency aware* |  |  |  |  |  |  |  |
| 100 ms | CS- – CS+ | 0.003 | 0.005 | 7580 | 0.720 | .471 | 0.181 |
| 1000 ms | CS- – CS+ | **-0.014** | **0.005** | **7580** | **-2.856** | **.008** | **-0.367** |
| 100 ms | HSF – LSF | 0.003 | 0.005 | 7580 | 0.555 | .579 | 0.148 |
| 1000 ms | HSF – LSF | **-0.010** | **0.005** | **7580** | **-1.973** | **.049** | **-0.354** |

Notes. *T*-statistics are evaluated for significance. A positive coefficient indicates higher ratings in the test condition than in the reference condition. Effects with $p \leq .05$ are marked in bold. Abbreviations: CS = conditioned stimulus, SE = standard error.
**P1: SCR subsample****Table S6. P1 Acquisition: Linear Mixed Model Fixed Effects (SCR subsample)**

### P1: SCR subsample

**Table S6.**
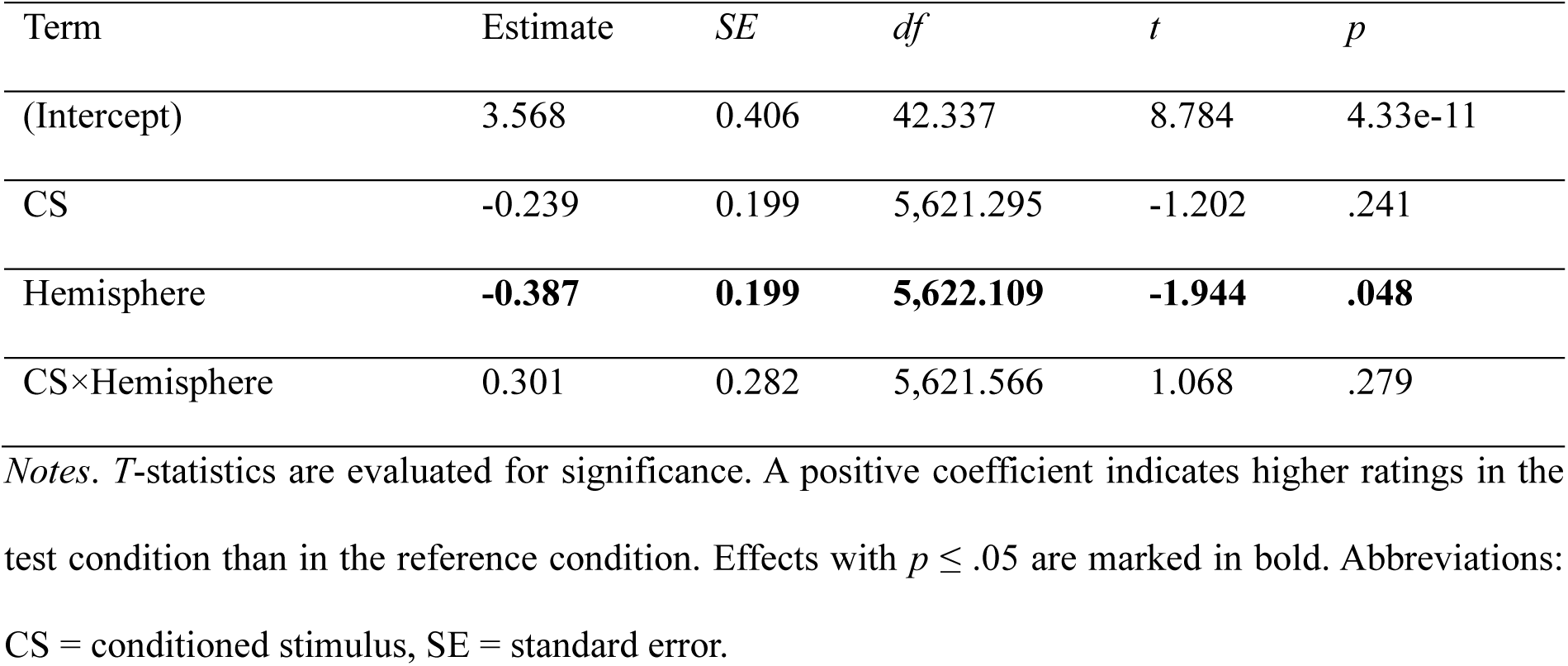
P1 Acquisition: Linear Mixed Model Fixed Effects (SCR subsample)

| Term | Estimate | SE | df | *t* | *p* |
| --- | --- | --- | --- | --- | --- |
| (Intercept) | 3.568 | 0.406 | 42.337 | 8.784 | 4.33e-11 |
| CS | -0.239 | 0.199 | 5,621.295 | -1.202 | .241 |
| Hemisphere | **-0.387** | **0.199** | **5,622.109** | **-1.944** | **.048** |
| CS×Hemisphere | 0.301 | 0.282 | 5,621.566 | 1.068 | .279 |

*Notes.* *T*-statistics are evaluated for significance. A positive coefficient indicates higher ratings in the test condition than in the reference condition. Effects with $p \leq .05$ are marked in bold. Abbreviations: CS = conditioned stimulus, SE = standard error.

**Table S7.**
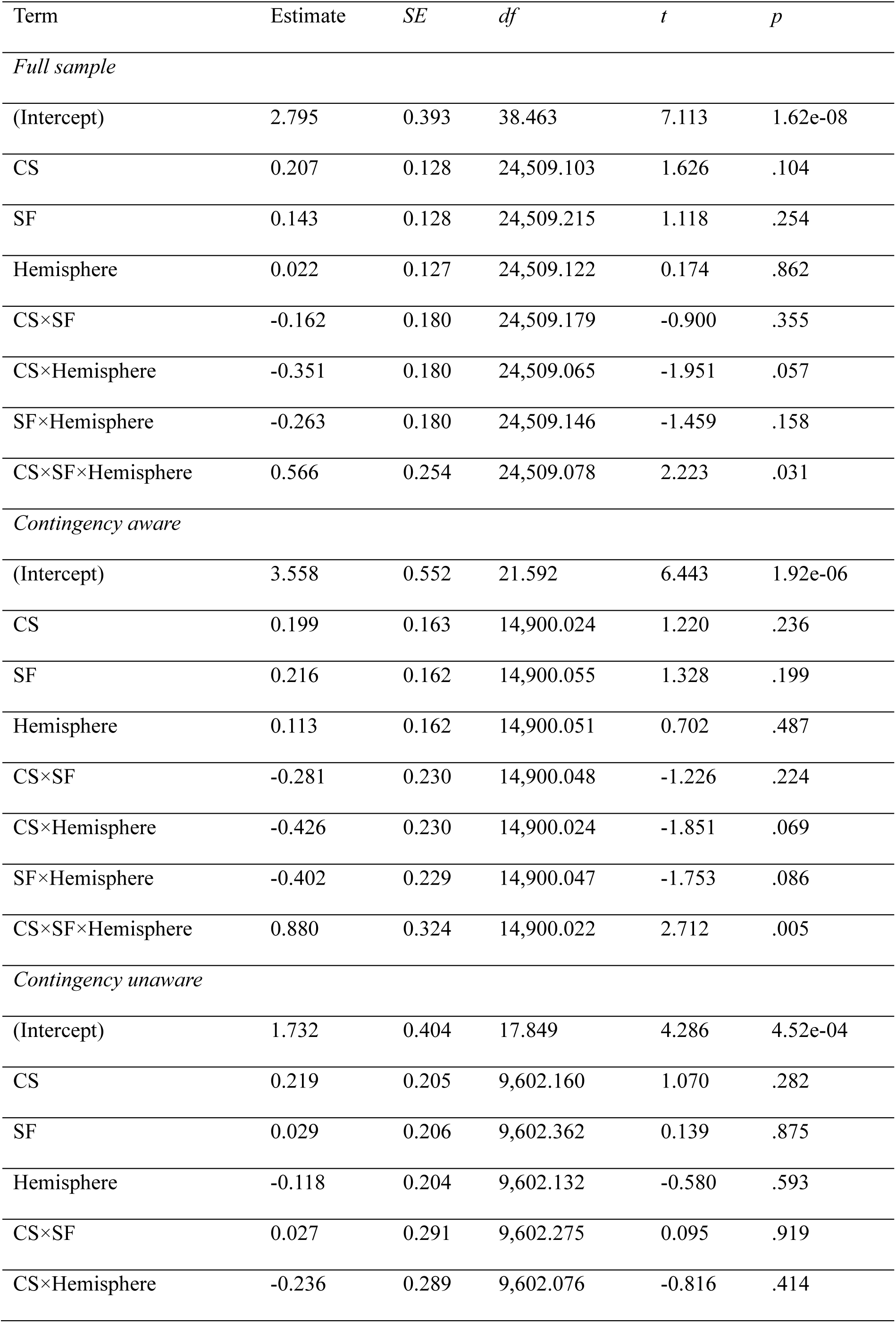
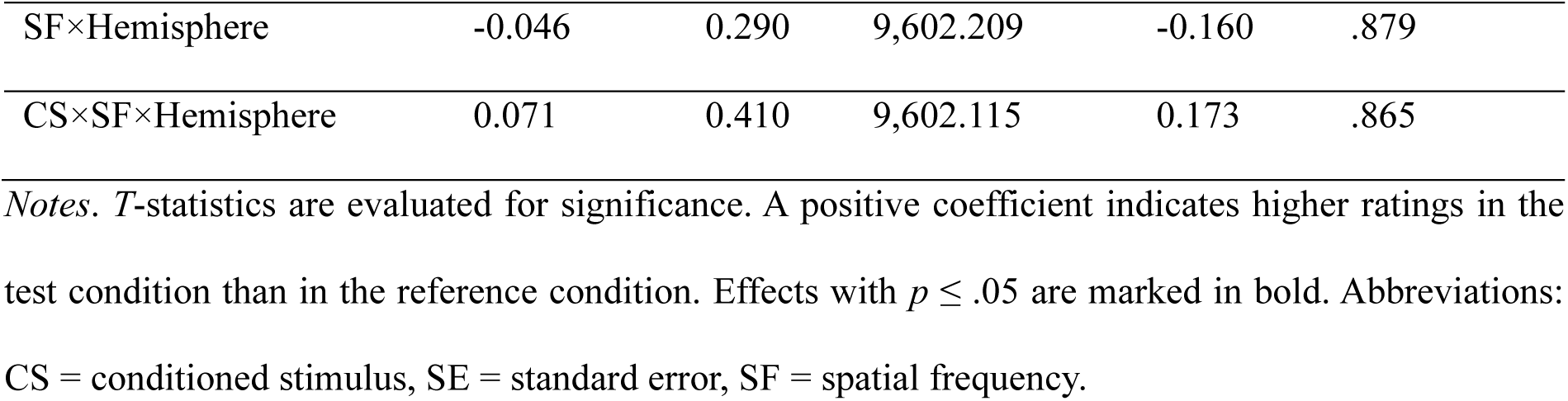
P1 Retrieval Linear Mixed Model Fixed Effects (SCR subsample)

**Table S8.**
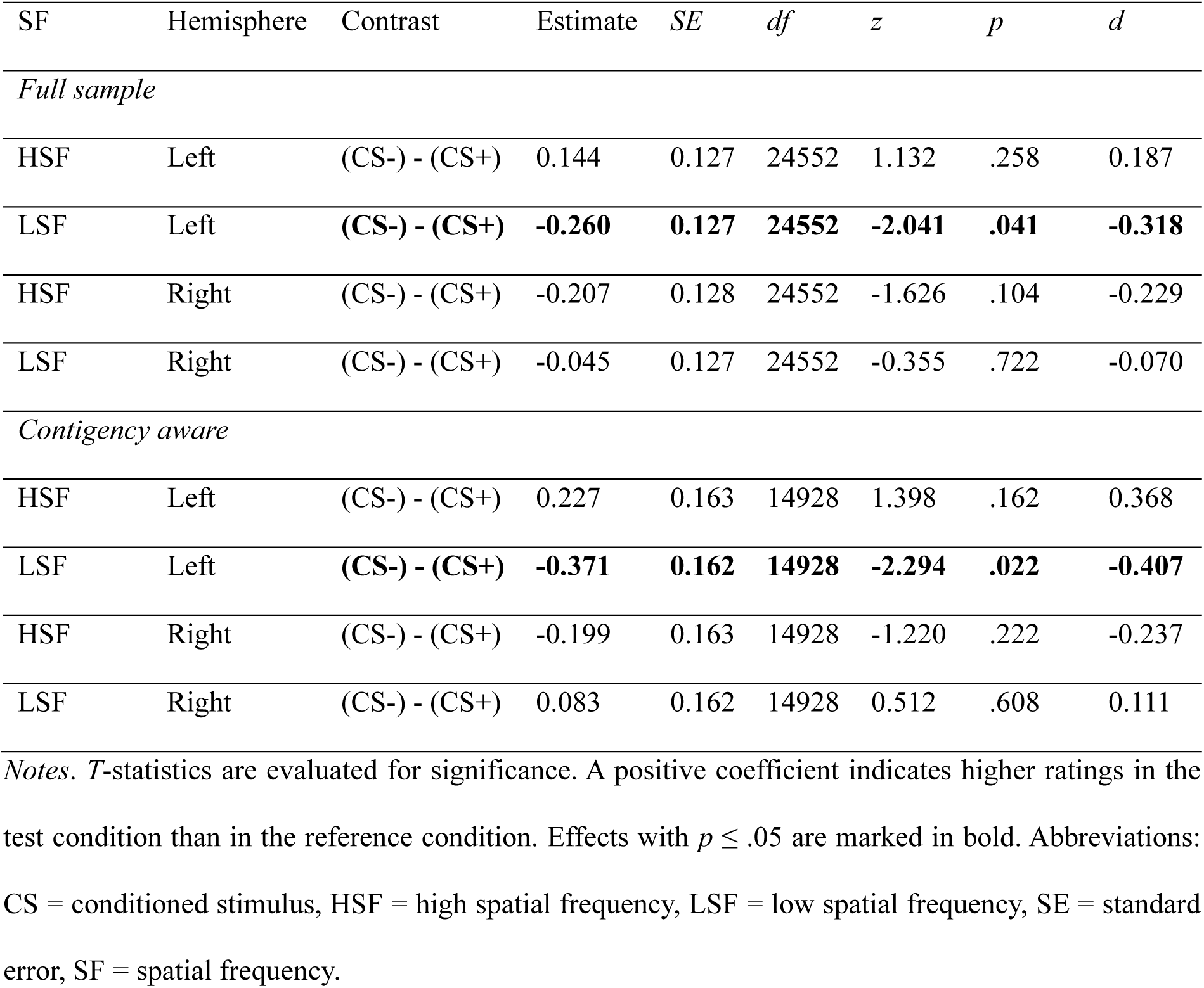
P1 Retrieval: Post-hoc CS Differences by SF × Hemisphere (SCR subsample)

### P1 – SCR association

**Table S9.** P1 – SCR association: Linear Mixed Model Fixed Effects.

**P1 – SCR association**1047 **Table S9. P1 – SCR association: Linear Mixed Model Fixed Effects**
| Term | Estimate | SE | df | t | p |
| --- | --- | --- | --- | --- | --- |
| *Full sample* |  |  |  |  |  |
| (Intercept) | **-0.13** | **0.05** | **44.007** | **-2.607** | **.012** |
| Trial | **1.77e-04** | **4.18e-05** | **9,866.211** | **4.222** | **2.45e-05** |
| EEG | **0.195** | **0.053** | **9,862.906** | **3.649** | **.035** |
| EEG×CS | -0.212 | 0.074 | 9,858.505 | -2.855 | .077 |
| EEG×SF | -0.181 | 0.075 | 9,858.793 | -2.421 | .112 |
| EEG×Hemisphere | -0.077 | 0.033 | 9,860.069 | -2.320 | .146 |
| EEG×CS×SF | 0.175 | 0.105 | 9,857.785 | 1.677 | .291 |
| EEG×CS×Hemisphere | 0.097 | 0.046 | 9,857.787 | 2.086 | .209 |
| EEG×SF×Hemisphere | 0.059 | 0.047 | 9,858.155 | 1.275 | .407 |
| EEG×CS×SF×Hemisphere | -0.055 | 0.065 | 9,857.907 | -0.839 | .625 |
| *Contingency aware* |  |  |  |  |  |
| (Intercept) | -0.132 | 0.07 | 24.286 | -1.903 | .069 |
| Trial | **2.03e-04** | **5.11e-05** | **6,927.802** | **3.970** | **7.26e-05** |
| EEG | 0.212 | 0.067 | 6,928.184 | 3.170 | .070 |
| EEG×CS | -0.207 | 0.095 | 6,925.042 | -2.182 | .205 |
| EEG×SF | -0.192 | 0.093 | 6,925.911 | -2.061 | .237 |
| EEG×Hemisphere | -0.079 | 0.041 | 6,926.271 | -1.930 | .249 |
| EEG×CS×SF | 0.16 | 0.132 | 6,924.612 | 1.213 | .477 |
| EEG×CS×Hemisphere | 0.095 | 0.059 | 6,924.782 | 1.624 | .350 |
| EEG×SF×Hemisphere | 0.055 | 0.057 | 6,925.435 | 0.961 | .579 |
| EEG×CS×SF×Hemisphere | -0.041 | 0.081 | 6,924.836 | -0.499 | .777 |
| *Contingency unaware* |  |  |  |  |  |
| (Intercept) | -0.121 | 0.068 | 19.967 | -1.779 | .091 |
| Trial | 1.10e-04 | 7.20e-05 | 2,929.595 | 1.530 | .065 |
| EEG | **0.171** | **0.088** | **2,924.742** | **1.946** | **.050** |
| EEG×CS | **-0.222** | **0.118** | **2,924.148** | **-1.878** | **.040** |
| EEG×SF | -0.174 | 0.123 | 2,923.709 | -1.405 | .134 |
| EEG×Hemisphere | -0.081 | 0.056 | 2,924.980 | -1.436 | .142 |
| EEG×CS×SF | 0.216 | 0.17 | 2,923.958 | 1.272 | .157 |
| EEG×CS×Hemisphere | 0.104 | 0.076 | 2,923.896 | 1.375 | .151 |
| EEG×SF×Hemisphere | 0.082 | 0.079 | 2,923.762 | 1.038 | .291 |
| EEG×CS×SF×Hemisphere | -0.098 | 0.109 | 2,923.978 | -0.902 | .320 |

**Table S10.**
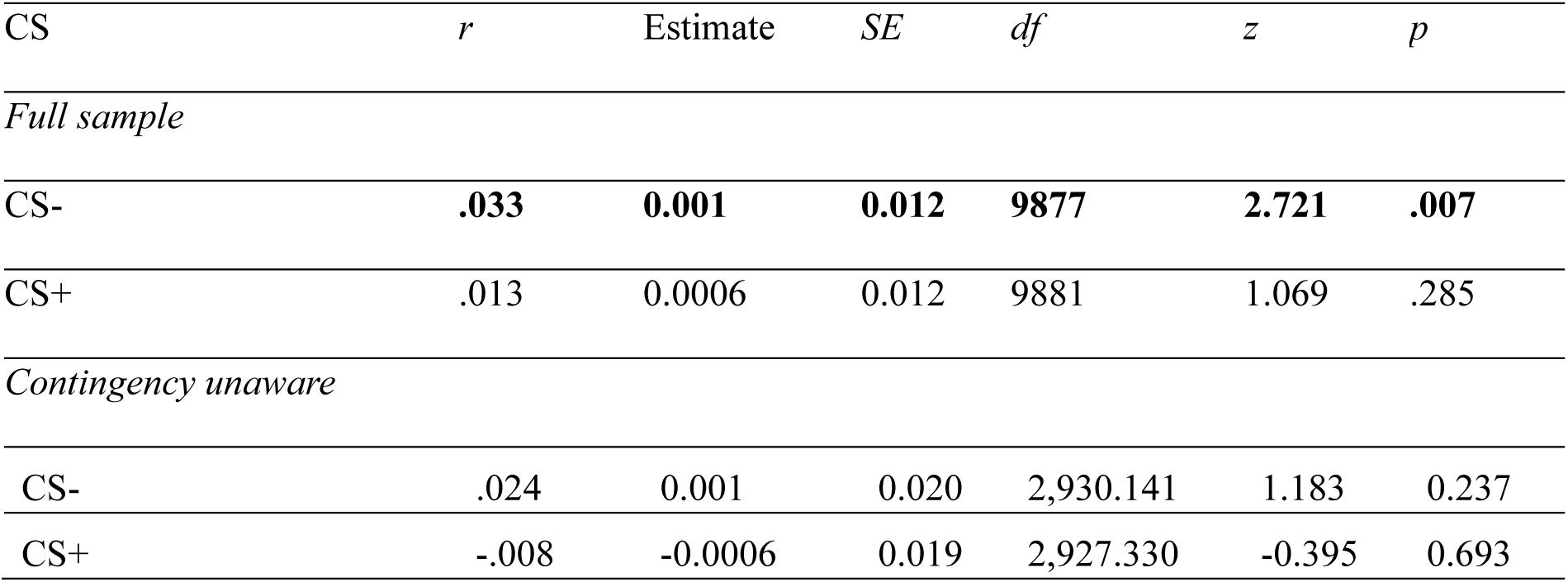
Post-hoc tests P1 – SCR association: Linear Mixed Model Fixed Effects.

| CS | *r* | Estimate | *SE* | *df* | *z* | *p* |
| --- | --- | --- | --- | --- | --- | --- |
| *Full sample* |  |  |  |  |  |  |
| CS- | **.033** | **0.001** | **0.012** | **9877** | **2.721** | **.007** |
| CS+ | .013 | 0.0006 | 0.012 | 9881 | 1.069 | .285 |
| *Contingency unaware* |  |  |  |  |  |  |
| CS- | .024 | 0.001 | 0.020 | 2,930.141 | 1.183 | 0.237 |
| CS+ | -.008 | -0.0006 | 0.019 | 2,927.330 | -0.395 | 0.693 |

## Notes

### Competing Interest Statement

The authors have declared no competing interest.

